# Isolation and characterization of two bacterial strains from textile effluents having Malachite Green dye degradation ability

**DOI:** 10.1101/2020.03.29.014274

**Authors:** Dipankar Chandra Roy, Md. Moinuddin Sheam, Md. Rockybul Hasan, Ananda Kumar Saha, Apurba Kumar Roy, Md. Enamul Haque, Md. Mizanur Rahman, Tang Swee-Seong, Sudhangshu Kumar Biswas

## Abstract

Water pollution from textile effluent is now one of the major issues all over the world. Malachite Green dye of the triphenylmethane group is a key component of textile effluents. This study aimed to isolate and identify potential Malachite Green dye degrading bacteria from textile effluents. Different growth and culture parameters such as temperature, pH, inoculum-size and dye concentration were optimized to perform the dye-degradation assay using different concentrations of Malachite Green dye in mineral salt medium. A photo-electric-colorimeter was used to measure the decolorizing activity of bacteria at different time intervals after aerobic incubation. Two competent bacterial strains of *Enterobacter* spp. (CV-S1 and CM-S1) were isolated from textile effluents showing potential degradation efficiency against Malachite Green dye. The RAPD analysis and 16S rRNA sequencing confirmed the genetical difference of the isolated strains *Enterobacter* sp. CV–S1 and *Enterobacter* sp. CM–S1. The two bacterial strains CV-S1 and CM-S1 showed complete Malachite Green dye degradation up to 15 mg/l under shaking condition with 5% (v/v) inoculums at pH 6.50 and temperature 35°C within 72 and 144 hours respectively. These findings indicate that the two potential bacterial strains can be used in large scale treatment of textile effluents in the future.

## 1. Introduction

Textile industries besides having a contribution to the development of the global economy, water pollution by textile effluents is one of the major concerned issues over the world. One million tons of synthetic dyes are produced each year worldwide and approximately 280,000 tons of these dyes are emitted to the industrial effluents annually [1][2]. Because of their non-degradable nature, these dyes inhibit the entrance sunlight in water and hamper the photosynthesis process, thus affecting the aquatic flora and fauna [3]. In addition, evaporation of these dyes and breathing them cause various allergic reactions and also hazardous for children [4]. In Brazil, several textile dyes that contaminate drinking water source have been reported to be carcinogenic [5].

Physio-chemical techniques, for example, coagulation-flocculation, adsorption, oxidation and membrane techniques have been used for decoloring of textile dyes during the past several decades. But these methods are associated with high sludge formation, high cost, generation of unwanted byproducts and secondary pollution [6]. Hence, there is an urgency of an eco-friendly, cost-effective and efficient approach of degrading textile dyes. Several microorganisms including bacteria, fungi and algae can be a potential alternative of removing polluting dyes [7]. Bioremediation of textile dyes by different microbial species have received much attention due to its cost-effectiveness, eco-friendly nature and public acceptability [8]. Isolation of textile dye degrading bacteria was started in the 1970s. Bacteria use these dyes as substrates and convert them into less complex metabolites by generating various oxidoreductive enzymes [9]. Recently different microbial species have been observed having the potentiality of degrading and decoloring textile dyes [10]–[15].

The present research aimed at isolating and characterizing Malachite Green (MG) dye degrading bacteria from the effluents of textile industry for possible use in the industrial process of bioremediation. Two bacterial strains *Enterobacter* CV-S1 and *Enterobacter* CM-S1 were identified for Malachite Green dye degrading capability.

## 2. Materials and methods

### 2.1 Sample collection

The untreated textile effluent samples, were collected in autoclaved plastic bottles from the drainage canal of two local textile industries in Kumarkhali, Kushtia, Bangladesh. Physical properties such as color, temperature and pH of the samples were recorded on the sites and were stored in the laboratory at 4°C within 12 hours of collection.

### 2.2. Dyes, chemicals and microbiological media

Triphenylmethane dye malachite green (MG) was procured from local market close to the thread dyeing plant of Kumarkhali, Kushtia, Bangladesh. Dye degrading mineral salt (MS) medium was prepared by adding the following components (in g/L): K_2_HPO_4_ (2), (NH_4_)SO_4_ (0.5), KH_2_PO_4_ (0.2) and MgSO_4_ (0.05); trace element (TE) Solution was prepared by adding the following components (in g/L): FeSO_4_ (0.4), MnSO_4_ (0.4), ZnSO_4_ (0.2), CuSO_4_ (0.04), KI (0.3), Na_2_MoO_4_ (0.05) and CoCl_2_ (0.04) and the components of enrichment medium was MS medium with glucose 0.1%, yeast extract 0.05%, peptone 0.5% and NaCl 0.5%. Furthermore, nutrient broth and nutrient agar medium were used for culture maintenance.

### 2.3 Isolation and screening of dye degrading bacteria

All samples (untreated textile effluents) were used for isolation of dye degrading bacterial cultures. *Enterobacter* sp. CV–S1 was isolated for Cristal Violet dye degradation as described in the previously published article (Roy et al., 2018) [7]. CV-S1 was further tested against MG dye. On the other hand, CM–S1 was isolated having both CV and MG dye degradation capability using the same techniques with few modifications.

### 2.4 Colony characteristics and microscopic morphology observation

After the isolation of two bacterial strains capable for significant dye degradation were the subject to study their various characteristics, such as study on colony characteristics, microscopic observation etc. Twenty four hour old cultures of the isolates were performed to study colony characteristics, Gram reaction and the cell morphology. The study of colony characteristics was determined according to Paul G. Engelkirk and Janet Duben-Engelkirk (2008) [16] and the cell morphology under microscope includes smear preparation and Gram’s staining was as described by WHO (2003) [17]

### 2.5 Growth characteristics determination

Bacterial growth characteristics was determined under different pH (6.00, 6.50, 7.00) and temperatures (30°C, 35°C and 40°C) as well as at the rotation of 120 rpm through growth curve analysis by turbidity measurements at different time intervals using photo-electric-colorimeter.

### 2.6 Sensitivity test to antibiotics

To identify drug resistance in bacteria, simple and well-standardized disc diffusion method was performed which was based on the measurement of zone of inhibition around each antibiotic disc [18]. Antibiotic of different classes, Ampicillin (10μg), Azithromycin (15μg), Bacitracin (10μg), Cephradine (30μg), Ceftriaxone (30μg), Doxycycline (30μg), Erythromycin (15μg), Neomycin (30μg), Sulphamethoxazole/Trimethoprim (25μg) and Tetracycline (30μg) were used to test antibiotic sensitivity.

### 2.7 Genomic DNA Extraction

The boiling method was used to extract the genomic DNA of the isolated bacteria. Concisely, a single colony from an overnight culture (at 37 °C) of LB agar plate was suspended into 30 µl of distilled water. The suspension was then boiled for 10 min at 100 °C. The mixture was instantly transferred into ice and cooled it for 5 min. The sample was centrifuged for 10 min at 13,000 × g and the supernatant, containing DNA, was used as the template for PCR amplification [19]

### 2.7.1 Characterization by RAPD analysis

In RAPD, synthetic (10 bases long) oligonucleotide primer is used to amplify genomic DNA under low annealing temperature [20]. The variable genes of the isolated bacterial strains were amplified using three RAPD primers: MT370563 (5’–TGCCGAGCTG–3’), MT370571 (5’–TGCGCCCTTC–3’) and MT370573 (5’–GTGAGGCGTC–3’). A total of 25μl reaction mixture for RAPD analysis contained ddH_2_O 14.75µl, MgCl_2_ (25mM) 2µl, buffer (10X) 2.5µl, dNTPs (10mM) 0.5µl, Taq DNA Polymerase (5u/µl) 0.25 µl, DNA template 1 µl and RAPD primer (10µM) 4µl. The Polymerase Chain Reaction was performed by Swift™ Minipro thermal cycler in the following steps: denaturing at 95 °C for 5 minutes, followed by 40 cycles of 40 seconds of denaturing at 95 °C, 60 seconds of annealing at 36 °C and 2 minutes of elongation at 72 °C with a final extension at 72 °C for 10 minutes.

### 2.7.2 Phylogenetic identification through 16S rRNA gene sequencing

16S rRNA genes were amplified using the bacteria-specific universal primers, a forward primer F27 (5’–AGAGTTTGATCCTGGCTCAG–3’; Tm: 61°C); and a reverse primer R1391 (5’–GACGGGCGGTGTGTRCA–3’; Tm: 67.4°C). The recipe of 25μl reaction mixture was: ddH_2_O 14.75µl, MgCl_2_ (25mM) 2µl, buffer (10X) 2.5µl, dNTPs (10mM) 0.5µl, Taq DNA Polymerase (5u/µl) 0.25 µl, DNA template 1 µl, forward primer (10µM) 2µl and reverse primer (10µM) 2µl. The thermal cycle for PCR amplification was same as mentioned in the section 2.7.1 except annealing temperature which was 65°C instead of 36°C. The amplified PCR product was purified from agarose gel by using AccuPrep® Gel Purification Kit (Bioneer corporation, Korea) in accordance with the directions of the manufacturer. The sequence was generated from the purified 16S rRNA gene using the DNA sequencer instrument (Model: Genetic Analyzer 3130). In order to identify the similarity in nucleotide sequences generated from the 16S ribosomal RNA amplification of the dye degrading bacteria was analyzed through NCBI BLAST (http://www.ncbi.nlm.nih.gov) by aligning the homologous sequences. Neighbor-joining method was used to determine evolutionary relationship by using online software: Muscle (v3.7), Gblocks (v0.91b), PhyML (v3.0 aLRT) and TreeDyn (v198.3) on the Phylogeny.fr platform [21], [22].

### 2.8 Assay of Malachite Green dye degradation

The rate of MG dye degradation was expressed as percentage. Using a photo electric colorimeter the degradation was determined through photo-electric-colorimeter by observing the decrease in absorbance at absorption maxima (λ max) where the degradation assay was determined by the following formula [23]. The concentration of dye in the supernatant was determined by absorbance at 660 nm.

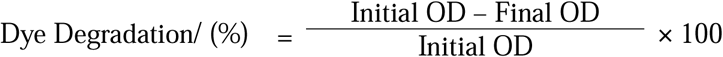

Various parameters like inoculum size, pH, temperature, and initial dye concentration were optimized as previously described by Roy *et al*. (2018) [7]. Briefly, a volume of 10ml solution containing MS medium enriched with trace element solution and MG dye was placed in 50ml test tube. The MG dye of 15, 30 and 50 mg/L concentrations were inoculated with the freshly grown 5% bacterial inoculums under aerobic shaking condition (120 rpm) at pH 6.50 and temperature 35°C. Each treatment was performed three times and the mean value was recorded for analysis.

## 3. Results and Discussion

### 3.1. Physical properties of textile effluents

Textile wastewater contains a high amount of unused dyes which create abnormal coloration [24]. Visually the physical appearance of the collected three textile effluent samples were black colored and one was turquoise blue. In addition, textile effluents have a wide PH and temperature range [25]. The pH determination of waste water is crucial as it is an important parameter that determines and influences the treatment process [26]. During collection the pH of tested sample was slightly acidic to neutral which was within the permissible limit of WHO and IS 10500:1991 [27]. Temperature affects various chemical and biological processes in water and is an important parameter [27]. Due to winter season the temperatures of the present collected samples were around 18°C.

### 3.2 Colony characteristics and microscopic morphology

The colony morphology or appearance of the colonies varies from one species to another and thus colony features serve as important clues in the identification of bacteria [16]. In this present investigation, though CV-S1 and CM-S1 were isolated with different samples, similar colony characteristics was observed as shown in Table 1. But according to the Figure 1, the gram negative rod shaped microscopic structures were significantly different in length which proved that both bacteria were microscopically different.

**Table 1:** Colony characteristics of MG dye degrading bacterial isolates on Nutrient agar

| Isolate | Color | Shape | Size | Opacity | Elevation | Surface texture | Stickiness |
| --- | --- | --- | --- | --- | --- | --- | --- |
| CV-S1 | White | Irregular | Large | Opaque | Flat | Rough | Yes |
| CM-S1 | White | Irregular | Large | Opaque | Flat | Rough | Yes |

**Figure 1:**
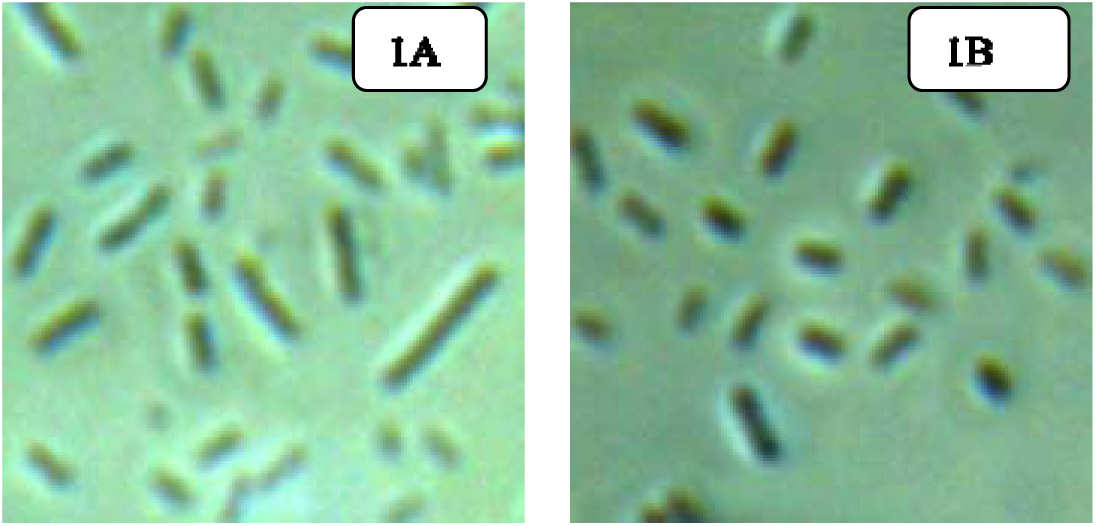
100× magnified microscopic morphology of MG dye degrading bacterial isolates: 1A: CV-S1; 1B: CM-S1

### 3.4 Growth characteristics

Growth pattern of bacteria can vary vastly species to species [28]. The growth curve of the microbial culture was analysed to study the population growth. The growth of microorganisms was plotted as the logarithm of the number of viable cells in the Y-axis and the incubation time in the X-axis. Growth curve (Figure 2) was obtained for two bacterial isolates, CV-S1 and CM-S1 showed growth variations at different temperature and pH gradients where maximum growth was observed at pH 6.50 and at temperature 35 °C.

**Figure 2:**
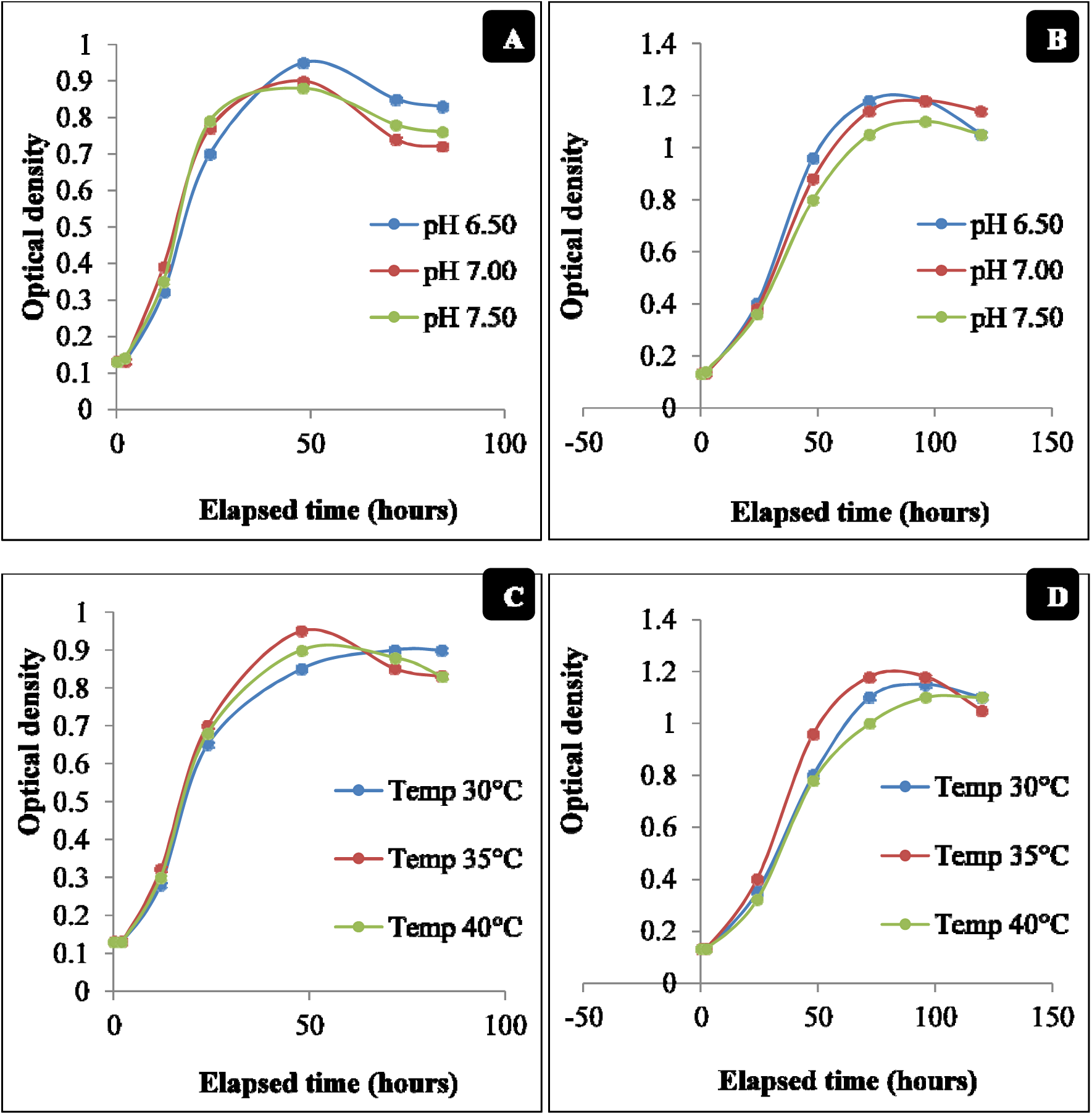
Effect of pH and temperature on bacterial growth: (A) Effect of pH on the growth of CV-S1;(B) Effect of pH on the growth of CM-S1; (C) Effect of temperature on the growth of CV-S1 and (D) Effect of temperature on the growth of CM-S1

### 3.5 Sensitivity to antibiotics

Globally, multi-drug resistant bacteria is an emerging public health crisis [29]. Wastewater treatment plant serves as a remarkable source of antibiotic resistant intestinal bacteria and antibiotic resistant genes, which may transfer to other environmental bacteria [30]. Hence, the determination of antibiotic resistance profile of selected bacteria is necessary for large scale implementation of these bacteria in treatment plants. Both CV-S1 and CM-S1 bacteria under this study based on the zone of inhibition were found sensitive to azithromycin, cefriaxone, sulphamethoxazole/trimethoprim and tetracycline, intermediate sensitive to ampicilin, doxycycline and neomycin and resistant to bacitracin, cephradine and erythromycin.

### 3.6 RAPD analysis

The two strains of MG dye degrading bacteria were characterized genetically by the Random Amplified Polymorphic DNA (RAPD) technique using the MT370563, MT370571 and MT370573 primers. Electrophoresis product of the amplified variable genes using 1.4% agarose gel showed different DNA banding patterns reveal that the two isolated bacterial strains CV–S1 and CMS–1 were genotypically different. Among three RAPD primers, primer MT370573 showed more variation than the primer MT370563 and MT370571 (Figure 3). Random amplification of polymorphic DNA (RAPD) is a PCR based method that is used to identify polymorphism in genomic DNA, thus useful in ascertaining dissimilarity in closely related bacterial strain [31].This molecular characterization of potential strains is helpful in identification and diversity analysis among bacterial isolates. [32] [33]

**Figure 3:**
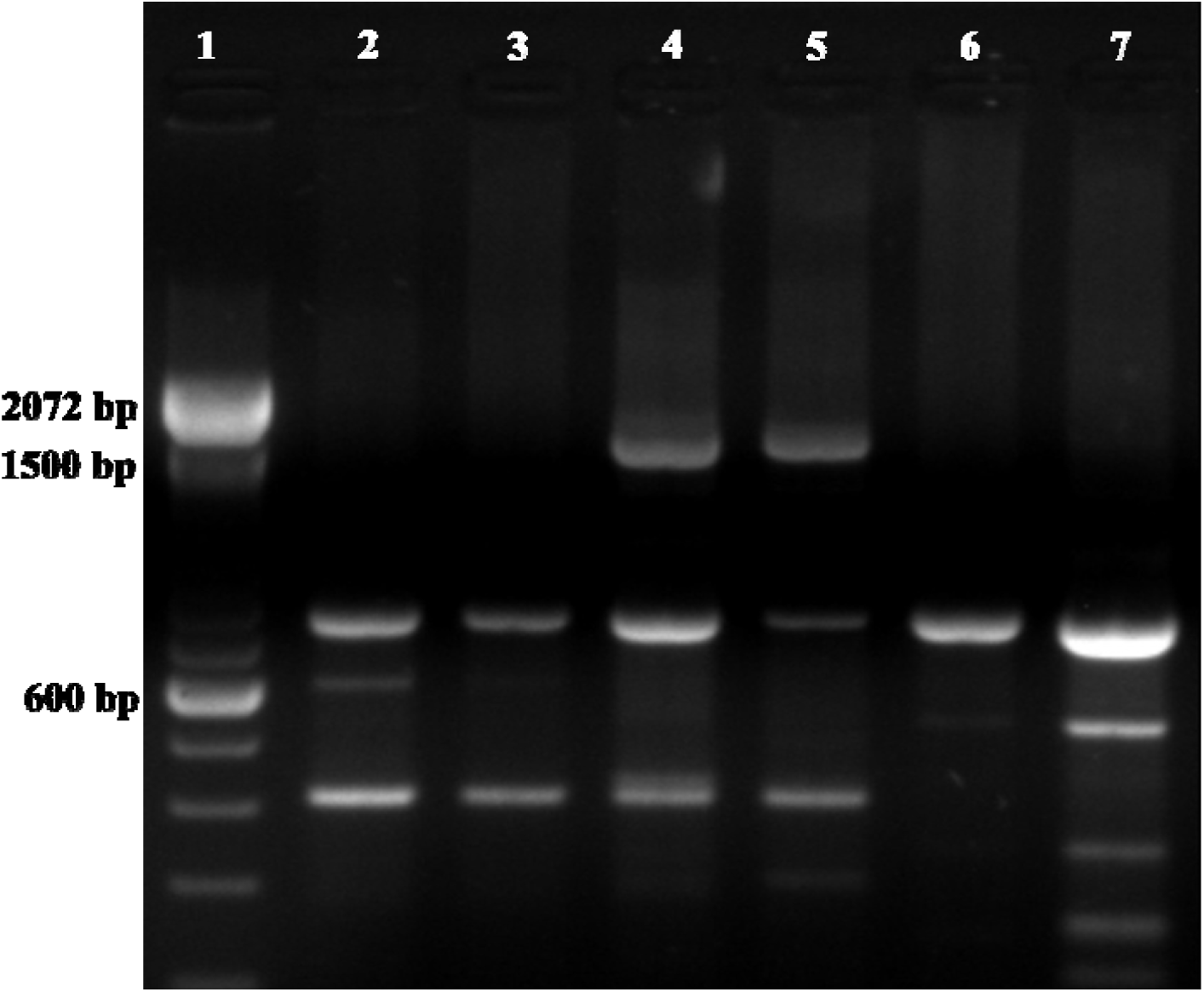
Observation of genotypic variation of two isolates using three RAPD primers: Lane 1, Marker; lane 2 and lane 3, strain CM-S1 and strain CV-S1 respectively with primer MT370563; lane 4 and lane 5, strain CM-S1 and strain CV-S1 respectively with primer MT370571 and lane 6 and lane 7, strain CM-S1 and strain CV-S1 respectively with primer MT37073

### 3.7 Phylogenetic identification of dye degrading bacteria

16S ribosomal RNA (rRNA) gene sequencing is a predominant strategy for bacterial indentification as well as phylogenetic relataionship analysis [34]. The tree exhibited highest identity 99% according to isolation source for CV–S1 was *Enterobacter* sp. HSL69 and for CM–S1 was *Enterobacter* sp. HSL99. In order to perform a better classification, phylogenetic tree was constructed. The evolutionary relationship of the dye degrading bacterial strains with other relevant bacteria were presented in the Figure 4 and their similarities found in the GenBank database in Table 2. The homology stipulated that the isolated strain CM–S1 was in the phylogenetic branch of the genus *Enterobacter*, and the strain CV–S1 constructed a new branch.

**Table 2:** Homology of the isolated bacterial strains CV–S1 and CM–S1 with other relevant bacterial strains found in the Gene Bank database

| Isolated Bacterial strain | Closely related bacteria | Accession no. | Identity (%) |
| --- | --- | --- | --- |
| CV-S1 | *Enterobacter cloacae* RU14 | KJ607595.1 | 99 |
|  | *Enterobacter cloacae* RJ04 | KC990807.1 | 99 |
|  | *Enterobacter* sp. HSL69 | HM461195.1 | 99 |
|  | *Enterobacter sacchari* SP1 | NR_118333.1 | 98 |
| CM-S1 | *Enterobacter cloacae* TPL2 | KJ470636.1 | 99 |
|  | *Enterobacter* sp. HSL99 | HM461229.1 | 99 |
|  | *Enterobacter* sp. HSL76B | HM461202.1 | 99 |
|  | *Enterobacter cloacae* RM20 | KJ607605.1 | 99 |

**Figure 4:**
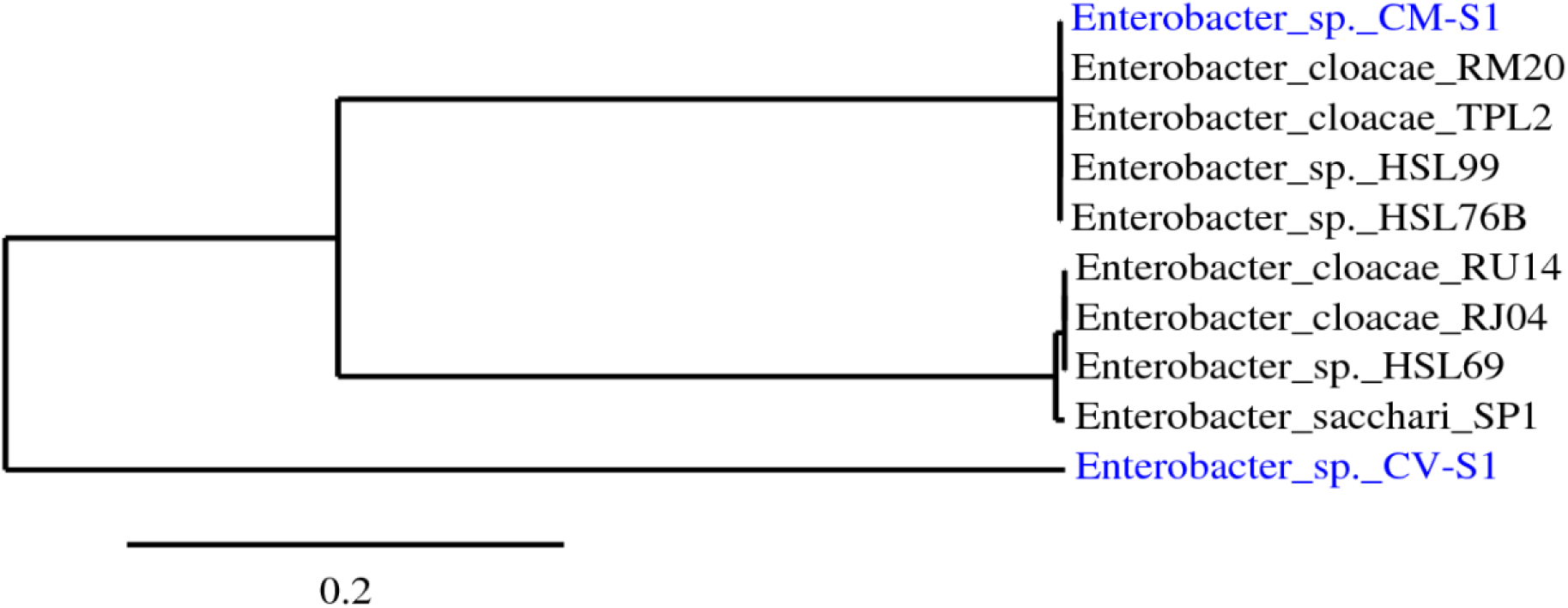
CM-S1 and CV-S1 were the isolated dye-degrading bacteria where the phylogenetic tree was reconstructed through maximum-homology method executed in the PhyML program (v3.0 aLRT) [21], [22].

Though identification of bacterial strains through 16S rRNA gene sequencing vary between genera and species, up to 90% bacterial strains at genus level and 86% at species level can be authentically identified by this process [35]. 98.65% similarity is recognized as threshold for species identification via 16S rRNA gene sequencing [36].Therefore, based on this consideration, the isolates were identified as *Enterobacter* sp. CV-S1 and *Enterobacter* sp. CM-S1. The newly constituted branch confirms that the identified strain CV-S1 is a new strain of the genus *Enterobacter*. [7]

### 3.8 Malachite green dye degradation

As pH 6.5 and temperature 35°C were optimized for dye degradation And at these defined pH and temperature, 5% inoculums of *Enterobacter* sp. CV-S1 was able to degrade completely up to 15 mg/L MG dye within 72 hours (Figure 5), whereas, 5% inoculums of *Enterobacter* sp. CM-S1 was able to degrade the same amount within 144 hours (Figure 6). Joshi and Mhatre **(2015)** also observed maximum MG dye degradation by *Enterobacter* sp. at pH 7.00 and temperature 37°C which was closely to this pH and temperature [37]. The degradation of malachite green was studied at various increasing concentration of dye i.e. from 15, 30 and 50 mg/l. It was found that the rate of degradation was decreased with increasing concentration of dye (Table 3 and Table 4) and similar trend was observed by Wanyonyi *et al*. (2018) [38]

**Table 3:**
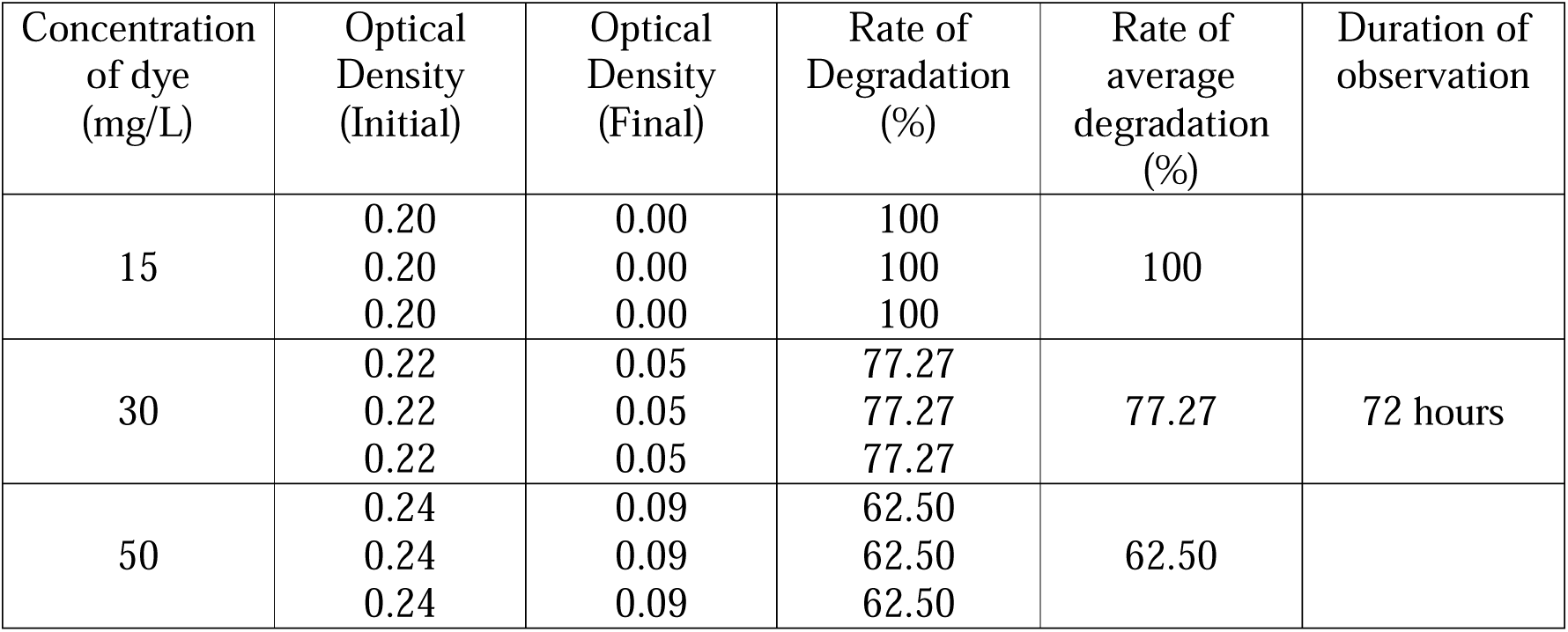
Different concentrations of MG dye degradation by *Enterobacter* sp. CV-S1

**Table 4:** Different concentrations of MG dye degradation by *Enterobacter* sp. CM-S1

| Concentration of dye (mg/L) | Optical Density (Initial) | Optical Density (Final) | Rate of Degradation (%) | Rate of average degradation (%) | Duration of observation |
| --- | --- | --- | --- | --- | --- |
| 15 | 0.20 | 0.00 | 100 | 100 |  |
|  | 0.20 | 0.00 | 100 |  |  |
|  | 0.20 | 0.00 | 100 |  |  |
| 30 | 0.22 | 0.09 | 59.09 | 59.09 | 144 hours |
|  | 0.22 | 0.09 | 59.09 |  |  |
|  | 0.22 | 0.09 | 59.09 |  |  |
| 50 | 0.24 | 0.11 | 54.17 | 54.17 |  |
|  | 0.24 | 0.11 | 54.17 |  |  |
|  | 0.24 | 0.11 | 54.17 |  |  |

**Figure 5:**
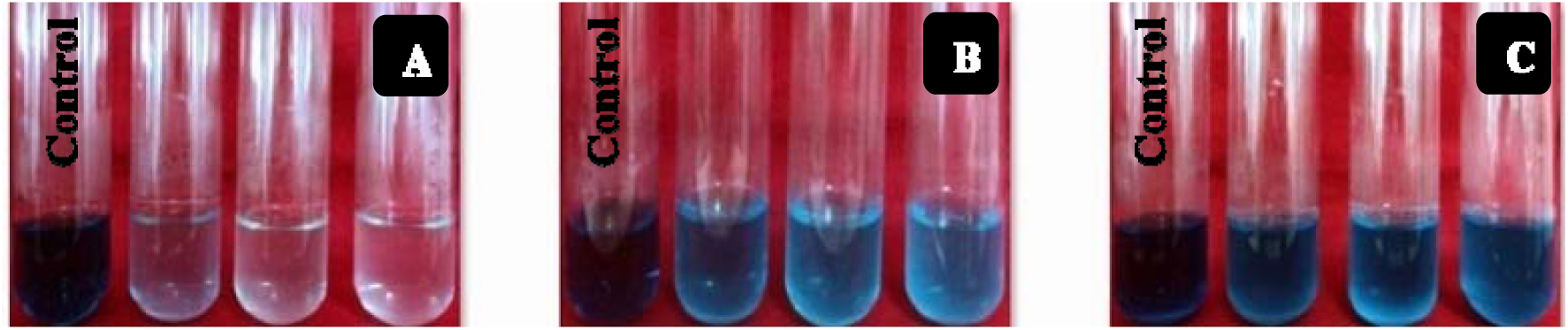
Degradation of Malachite green dye by *Enterobacter* sp. CV-S1: **A**, 15 mg/L; **B**, 30 mg/L; **C**, 50 mg/L

**Figure 6:**
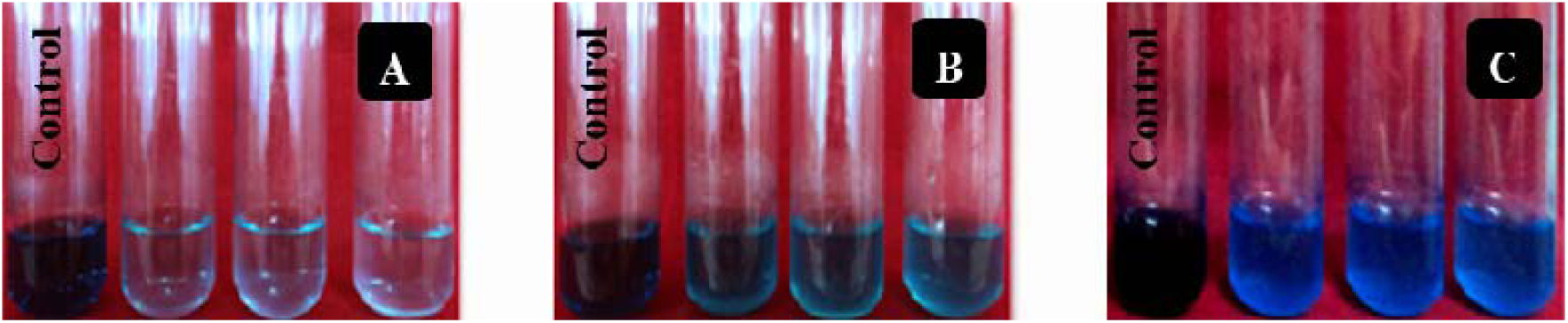
Degradation of Malachite green dye by *Enterobacter* sp. CM-S1: **A**, 15 mg/L; **B**, 30 mg/L; **C**, 50 mg/L

Another interesting finding is that during MG dye degradation no extra carbon and protein source was added in MS medium i.e. both bacteria had used MG dye as a carbon sources. But most of researches carbon or protein-enriched media were used for MG dye degradation by bacteria [39]–[42]. Indeed, those organisms are the real degrading organisms which can utilize MG dye as a sole carbon source and their degrading capacity in the presence of a trace amount of carbon source is dramatically accelerated. In a study Ali et al (2009), observed MG dye degradation for 6 days by two fungal strains *Aspergillus flavus* and *Alternaria solani* up to 30µM (≈10.95mg/L) when MG was a sole source of carbon but degradation had increased up to 50µM (≈18.25mg/L) when extra carbon source was added in the medium [43]. Hence, *Enterobacter* sp. CV-S1 and *Enterobacter* sp. CM-S1 may be more effective industrially as they utilized up to 15mg/L MG dye as a carbon source.

## 4 Conclusion

Although microbial degradation/decolorization of textile effluent is a challenging process, some microbes recently have been explored as the potential scavengers in textile dye and textile effluent treatment. In current study, two gram-negative rod shaped potential bacterial strains *Enterobacter* sp. CV-S1 and *Enterobacter* sp. CM-S1 were identified having MG dye degradation capacity. As these isolates were able to degrade significant amount of MG dye without providing any carbon source, adding desirable carbon source they can be used successfully in the commercial treatment of textile effluents. These observations reveal that the two potential bacterial strains could be used in water treatment plant in near future.

